# Flow-field inference from neural data using deep recurrent networks

**DOI:** 10.1101/2023.11.14.567136

**Authors:** Timothy Doyeon Kim, Thomas Zhihao Luo, Tankut Can, Kamesh Krishnamurthy, Jonathan W. Pillow, Carlos D. Brody

## Abstract

Computations involved in processes such as decision-making, working memory, and motor control are thought to emerge from the dynamics governing the collective activity of neurons in large populations. But the estimation of these dynamics remains a significant challenge. Here we introduce Flow-field Inference from Neural Data using deep Recurrent networks (FINDR), an unsupervised deep learning method that can infer low-dimensional nonlinear stochastic dynamics underlying neural population activity. Using population spike train data from frontal brain regions of rats performing an auditory decision-making task, we demonstrate that FINDR outperforms existing methods in capturing the heterogeneous responses of individual neurons. We further show that FINDR can discover interpretable low-dimensional dynamics when it is trained to disentangle task-relevant and irrelevant components of the neural population activity. Importantly, the low-dimensional nature of the learned dynamics allows for explicit visualization of flow fields and attractor structures. We suggest FINDR as a powerful method for revealing the low-dimensional task-relevant dynamics of neural populations and their associated computations.

## 1. Introduction

One of the major challenges in systems neuroscience is in identifying the right level of abstraction to describe how a neural system functions, and bridging such a description to both the cellular-level implementation and behavior. In one approach, we start with the computational task that the neural system has to solve, and either hand-build (Hopfield, 1982; Gerstner & van Hemmen, 1992; Wang, 2002) or train a network of model neurons (Sussillo & Barak, 2013; Yang et al., 2019; Dubreuil et al., 2022; Driscoll et al., 2022) to solve this task. While these networks provide insights into how individual model neurons could work together to solve a particular task, these model neurons are often not directly constrained to capture the heterogeneous responses observed in real neurons. It is therefore possible that the mechanisms used by these networks to solve a task do not fully reflect the mechanisms used by real neural populations in the brain.

In another approach, we start with the neural population activity measured from an animal performing a computational task, and attempt to infer latent representations, or factors, that are relevant to the task computations (Cunningham & Yu, 2014). The dynamics of these representations (i.e., how they evolve over time) are thought to mediate the computations performed by neural populations (Vyas et al., 2020; Duncker & Sahani, 2021), and unsupervised methods have been developed to infer these dynamics from neural population activity. Currently available methods make simplifying assumptions on the dynamics to facilitate inference. For example, dynamics are assumed to be autonomous (Duncker et al., 2019), linear (Macke et al., 2011; Gao et al., 2016), switching linear (Linderman et al., 2017; Nassar et al., 2019; Zoltowski et al., 2020), deterministic except at specific time points (Pandarinath et al., 2018; Kim et al., 2021; Keshtkaran et al., 2022), or high-dimensional (Pandarinath et al., 2018; Keshtkaran et al., 2022). However, the assumptions on dynamics made by these inference methods may not necessarily align with the dynamics in real neural populations. Moreover, most currently available methods, applied naively, do not distinguish between task-relevant and -irrelevant dynamics. The complex response patterns observed in real neurons may in part be due to their mixed selectivity to task-related and -unrelated variables (Rigotti et al., 2013), and the methods we use should separate the task-relevant and -irrelevant components in the neural population responses.

To address these gaps, we propose a novel method called FINDR (Flow-field Inference from Neural Data using deep Recurrent networks) and present an overview of the method in Sections 2.1–2.3. The FINDR method builds on the existing methods for latent dynamics inference in two major ways. First, leveraging the flexibility of a recently introduced class of dynamical models called neural stochastic differential equations (nSDE; Li et al. (2020); Kidger et al. (2021a;b)), FINDR learns nonlinear stochastic latent dynamics. This allows FINDR to capture the heterogeneous responses of individual neurons in large populations. We demonstrate in Section 2.5 that FINDR outperforms existing methods in reconstructing the individual neural responses from frontal cortical regions involved in decision-making. Second, we take measures to improve the interpretability of the FINDR-inferred latent dynamics. We constrain the learned dynamics to lie on a low-dimensional latent space and infer how the given external inputs to the system influence the dynamics. Furthermore, we infer the task-relevant and -irrelevant dynamics separately, so that we can focus only on the dynamics of the population that are relevant to the task computations. In Section 2.4, we show that FINDR can infer interpretable low-dimensional attractor structures from spike trains generated by synthetic neural populations that memorize continuous quantities. FINDR takes the continuous quantities used by the populations as its inputs and uses appropriate attractor structures to maintain the memory of these quantities. We also demonstrate in Section 2.5 that FINDR can express neural population activity in terms of latent representations of dimensions lower than existing methods. Because FINDR can represent neural population activity in low dimensions, even as low as two or three dimensions, we can explicitly visualize the flow field (or the velocity vector field) underlying neural population activity. In Section 2.6, we show that when we visualize the vector field formed by the frontal cortical neural population during decision-making, we see two attractors, with each of them representing a choice alternative.

## 2. Results

### 2.1. Goal of FINDR

Given some population spike train data from an animal performing an experimental task, we want to identify a low-dimensional latent space ℝ^*L*^ that is embedded in the neural population state space 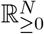, with the mapping 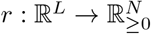 given by

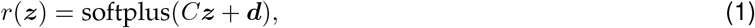

where *C* is a semi-orthogonal loading matrix and ***d*** is the time-varying bias. Here, the softplus nonlinearity prevents the firing rates from being negative. Instead of Equation (1), we could have used a more general mapping *r*(***z***) = softplus(Ψ_*κ*_(***z***)), where Ψ is a differentiable map with parameters *κ*, and could, for example, be a deep neural network. However, for simplicity and for interpretability (see Section 3), we confine our map to be affine. Notably, our semi-orthogonal *C* makes the distance and angle in the latent space ℝ^*L*^ equivalent to the distance and angle in the inverse-softplus rate space ℝ^*N*^ (see Section 4.2). The observed population spike trains ***y*** are modeled by non-homogeneous Poisson processes with rates 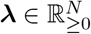 conditioned on the latent variable ***z*** ∈ ℝ^*L*^:

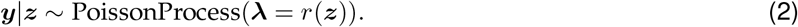

The instantaneous firing rates ***λ*** = *r*(***z***) change over time *t* according to a stochastic differential equation (SDE) of the form

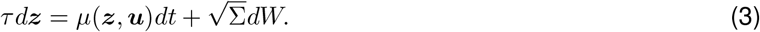

Here, *W* is a Wiener process, *τ* is a fixed time constant of the SDE, and ***u*** is the external input to the system, ypically a set of task variables that the experimenter has control over. The goal of FINDR is to find the drift _79_ ction *μ*, the noise covariance Σ, and the mapping *r* that best capture the observed population spike trains ***y***.

### 2.2. Gated Neural Stochastic Differential Equations

In FINDR, the drift function *μ* in Equation (3) is approximated by

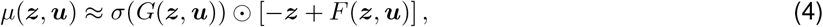

where *σ* is the sigmoid function that acts element-wise, and *F* and *G* are feedforward neural networks (FNNs). A model parameterized by *d****z****/dt* = *σ*(*G*(***z, u***)) ⊙ [− ***z*** + *F* (***z, u***)] is known as the gated neural ordinary differential equation (gnODE) (Kim et al., 2023). The gnODE is a recurrently connected network model that has been shown to outperform recurrent neural networks (RNNs) that achieve near state-of-the-art (Rusch et al., 2021) on various tasks, particularly when *L* is low (Kim et al., 2023). We use a novel variant of the gnODE, the gated neural stochastic differential equation (gnSDE), that also models noise in the dynamics by learning the covariance matrix Σ (which we assume is diagonal for simplicity, but can in principle be any symmetric and positive semi-definite matrix) along with the parameters of the gated FNNs *F* and *G*.

Noise in the latent dynamics, i.e., 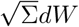 in Equation (3), can be a crucial component in modeling neural computation. As an example, in perceptual decision-making, we observe that even when an animal is presented with identical stimuli, the animal’s choice behavior can vary from trial to trial. If we are given spiking observations from a neural population that represents the animal’s choice, we can model variability in the animal’s behavior with noise in the dynamics of the neural population. Poisson noise at the level of spiking observation in Equation (2) may not be sufficient to model this variability.

### 2.3. FINDR architecture and inference procedure

At the implementational level, we work with the discretized analogs of the continuous variables in Equations (1-3) with step size Δ*t* = 0.01s. Going forward, we denote ***z***_*k*_ = ***z***(*k*Δ*t*), ***u***_*k*_ = ***u***(*k*Δ*t*), ***r***_*k*_ = *r*(***z***(*k*Δ*t*)), ***d***_*k*_ = ***d***(*k*Δ*t*), ***y***_*k*_ = ***y***(*k*Δ*t*), with *k* = 1, 2, 3, …, *T*, where *T* is the total number of steps taken in a trial.

FINDR is implemented as a variational autoencoder (VAE; Kingma & Welling (2014)) with a sequential structure (e.g., Chung et al. (2016); Krishnan et al. (2016); Pandarinath et al. (2018)) shown in Figure 1. It minimizes two objectives: one for neural activity reconstruction (Figure 1a) and the other for flow-field (i.e., velocity vector-field) inference (Figure 1b-c). For the first objective, FINDR receives some population spike trains ***y***_1:*T*_ and task-relevant external inputs ***u***_1:*T*_, and outputs the reconstructed population spike trains ***ŷ***_1:*T*_ from the latent trajectory ***z***_1:*T*_ and bias ***d***_1:*T*_ (Figure 1a). To make the learned latent ***z***_1:*T*_ maximally relevant to the task, we first estimate, for each neuron, its baseline fluctuations in the firing rate across trials independent of task conditions using a set of radial basis functions. We use the estimated quantities as our time-varying bias ***d***_1:*T*_, and call ***d***_1:*T*_ the task-irrelevant baseline inputs (see Section 4.1.1 for details). With the baseline input ***d***_1:*T*_ fixed, we then train the rest of the model. In inferring the latent ***z***_*k*_ for each time point *k*, FINDR makes use of the representation ***e***_*k*_ learned by a bidirectional GRU (Cho et al., 2014), which has information about the entire trajectory of the population spike trains ***y***_1:*T*_ and external inputs ***u***_1:*T*_ . The learned ***z***_*k*_ then gets transformed to 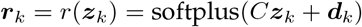, following Equation (1). We reconstruct the spiking activity ***y***_*k*_ with

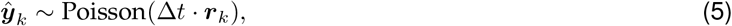

which is the discretized form of Equation (2). The reproduced activity ***ŷ***_*k*_ will become more similar to the observed activity ***y***_*k*_ as we minimize the reconstruction objective

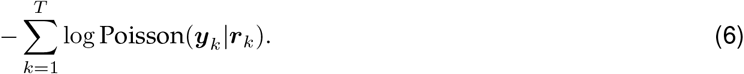

For the second objective, FINDR must learn how the task-relevant latent ***z***_*k*_ evolves over time. This is done by a gnSDE discretized with the Euler-Maruyama method (Figure 1b):

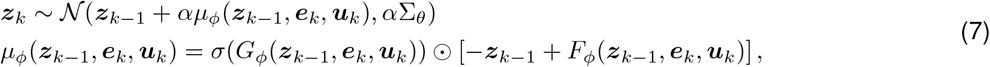

where Σ_*θ*_ is a diagonal matrix, *α* = Δ*t/τ*, and *F*_*ϕ*_ and *G*_*ϕ*_ are FNNs. We call *μ*_*ϕ*_ the “posterior drift network”. Sampling ***z***_1:*T*_ from Equation (7) is equivalent to the sampling procedure from the variational approximation of the posterior *q*_*ϕ*_(***z***_1:*T*_ |***y***_1:*T*_, ***u***_1:*T*_, ***d***_1:*T*_) of the true posterior *p*_*θ*_(***z***_1:*T*_ |***y***_1:*T*_, ***u***_1:*T*_, ***d***_1:*T*_) in VAEs (Kingma & Welling (2014); see Section 4.1.2 for details). Note that the posterior drift network *μ*_*ϕ*_(***z***_*k*−1_, ***e***_*k*_, ***u***_*k*_) outputs the velocity *ż* _*k*_ of ***z***_*k*_ at time point *k*, given ***e***_*k*_ and ***u***_*k*_. However, to infer the SDE in Equation (3), we must know what the velocity ***ż***_*k*_ is at time point *k*, given only ***u***_*k*_:

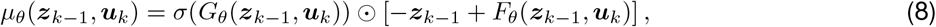

where *F*_*θ*_ and *G*_*θ*_ are FNNs that are different from *F*_*ϕ*_ and *G*_*ϕ*_. We call *μ*_*θ*_ the “prior drift network”. To match the velocity *ż*_*k*_ learned by the posterior drift network *μ*_*ϕ*_(***z***_*k*−1_, ***e***_*k*_, ***u***_*k*_) and the prior drift network *μ*_*θ*_(***z***_*k*−1_, ***u***_*k*_), we minimize the squared Mahalanobis distance *D*_*M*_, which is equivalent to a weighted mean squared error (MSE) in our case (Figure 1c):

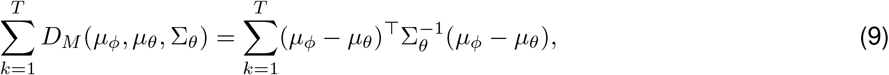

where the arguments of the drift networks *μ*_*ϕ*_ and *μ*_*θ*_ are (***z***_*k*−1_, ***e***_*k*_, ***u***_*k*_) and (***z***_*k*−1_, ***u***_*k*_), respectively. It can be shown that *D*_*M*_ is equivalent to the Kullback-Leibler divergence *D*_KL_ between the variational posterior *q*_*ϕ*_(***z***_1:*T*_ |***y***_1:*T*_, ***u***_1:*T*_, ***d***_1:*T*_) and the prior *p*_*θ*_(***z***_1:*T*_ |***u***_1:*T*_) (Kingma & Welling (2014); see Section 4.1.2 for details).

**Figure 1.**
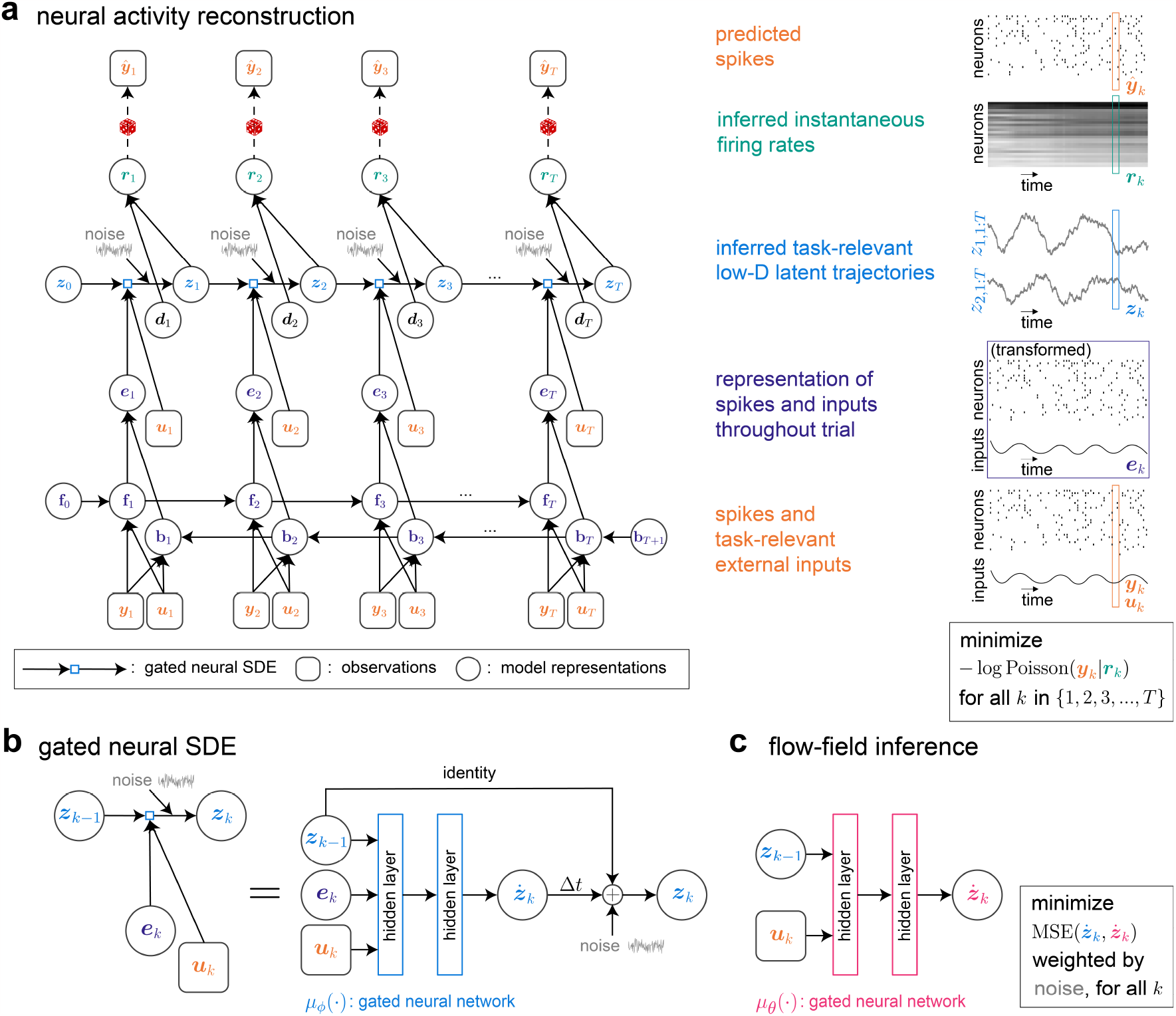
Directed graphs of FINDR. FINDR reconstructs neural population activity from low-dimensional latent stochastic dynamics. **a**, Reconstruction is done by first learning a representation that summarizes the observed population spike trains ***y***_1:*T*_ and task-relevant external input ***u***_1:*T*_ into the variable ***e***_*k*_ = [**f**_*k*_; **b**_*k*_], using a bidirectional GRU that runs forward (**f**_*k*_ = GRU(**f**_*k−*1_, ***y***_*k*_, ***u***_*k*_)) and backward (**b**_*k*_ = GRU(**b**_*k*+1_, ***y***_*k*_, ***u***_*k*_)) in time. We infer the task-relevant low-dimensional latent ***z***_*k*_ at each time point *k* by using ***e***_*k*_ and ***u***_*k*_. The task-relevant latent ***z***_*k*_ and the task-irrelevant baseline inputs ***d***_*k*_ are used to infer the instantaneous firing rates of individual neurons ***r***_*k*_. The initial condition ***z***_0_ is set to be **0**. Parameters **f**_0_ and **b**_*T* +1_ are trained. We minimize the Poisson negative log-likelihood of observing spikes ***y***_*k*_ given the firing rates ***r***_*k*_. **b**, The evolution of the latent ***z***_*k*_ over time is learned by a gated neural SDE (gnSDE). The gnSDE is parametrized by a gated FNN *μ*_*ϕ*_ (the posterior drift network), which outputs the velocity ***ż***_*k*_ given ***z***_*k−*1_, ***e***_*k*_, and ***u***_*k*_. **c**, Using a separate gated FNN *μ*_*θ*_ (the prior drift network), we learn how the velocity ***ż***_*k*_ depends on ***z***_*k−*1_ and ***u***_*k*_.

Combining Equation (6) and Equation (9), we get the following objective ℒ:

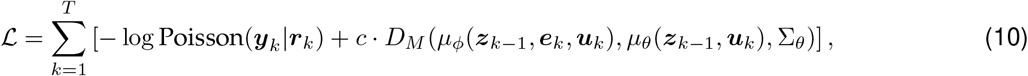

where the weight coefficient *c* determines the relative importance of the inferred velocity vector field and the reconstructed neural activity.

*c* is an important hyperparameter in Equation (10) that determines the trade-off between the accuracy of the reconstructed neural activity and the discovered vector field. When *c* is too low, the reconstructed neural activity may be accurate, but it becomes unlikely that the inferred latent trajectory ***z***_1:*T*_ is generated from the vector field ***ż***= *μ*_*θ*_(***z, u***). If *c* is too high, the inferred latent trajectory ***z***_1:*T*_ becomes more irrelevant to the observed neural activity, but it becomes highly likely that ***z***_1:*T*_ is generated from our inferred vector field ***ż***= *μ*_*θ*_(***z, u***). When *c* = *α/*2 = Δ*t/*2*τ*, the objective *ℒ* in Equation (10) can be shown to be an upper bound of the marginal negative log-likelihood of observing the spiking data ***y***_1:*T*_ given the external inputs ***u***_1:*T*_ and separately computed baselines ***d***_1:*T*_, i.e., − log *p*(***y***_1:*T*_ |***u***_1:*T*_, ***d***_1:*T*_) (Kingma & Welling (2014); see Section 4.1.2 for details). Computing the marginal negative log-likelihood − log *p*(***y***_1:*T*_ |***u***_1:*T*_, ***d***_1:*T*_) can be computationally expensive, and we therefore use *ℒ* as our objective for efficient training of our model. It has been observed that using *c > α/*2 can help arrive at interpretable latent representations (Higgins et al., 2017; Burgess et al., 2018). We let *c* = *α* = Δ*t/τ* to put slightly more weight on the vector-field inference at the cost of less accurate reconstruction of neural activity. However, as we will show in Section 2.5, we find that under many conditions, FINDR outperforms existing methods in reconstructing neural population activity.

One may ask why we train two separate networks, the posterior drift network *μ*_*ϕ*_ and the prior drift network *μ*_*θ*_, instead of letting the posterior drift network *μ*_*ϕ*_ be equivalent to the prior drift network *μ*_*θ*_. This reduces the number of model parameters to train, and can potentially be helpful in the data-limited regime. However, this also means that FINDR should reconstruct spiking activity ***ŷ***_*k*_ based only on ***u***_1:*k*_. This tends to hurt the reconstruction performance. In other words, interpreting FINDR within the Bayesian framework, the variational approximation *q*_*ϕ*_(***z***_1:*T*_ |***y***_1:*T*_, ***u***_1:*T*_, ***d***_1:*T*_) of the true posterior *p*_*θ*_(***z***_1:*T*_ |***y***_1:*T*_, ***u***_1:*T*_, ***d***_1:*T*_) is parametrized by the posterior drift network *μ*_*ϕ*_ and the bidirectional GRUs – we compute the approximation *q*_*ϕ*_ rather than *p*_*θ*_(***z***_1:*T*_| ***y***_1:*T*_, ***u***_1:*T*_, ***d***_1:*T*_) because computing the latter is computationally expensive. If the variational approximation *q*_*ϕ*_ is set to be equivalent to the prior *p*_*θ*_(***z***_1:*T*_ |***u***_1:*T*_) (i.e., *μ*_*ϕ*_ = *μ*_*θ*_), then L will not be a tight upper bound on the marginal negative log-likelihood − log *p*(***y***_1:*T*_| ***u***_1:*T*_, ***d***_1:*T*_) (Kingma & Welling, 2014). Therefore, we want the posterior drift network *μ*_*ϕ*_ and the prior drift network *μ*_*θ*_ to be different networks (see Section 4.1.2 for details).

We train FINDR by computing the gradient of *ℒ* in Equation (10) with respect to the parameters of the bidirectional GRU, *F*_*ϕ*_, *G*_*ϕ*_, *F*_*θ*_, *G*_*θ*_, Σ_*θ*_, and *C*, and doing mini-batch gradient descent with warm restarts (Loshchilov & Hutter (2017); see Section 4.1.5 for details). The gradient is obtained by backprogation through time (BPTT).

### 2.4. FINDR can accurately discover latent continuous attractors in synthetic neural populations

To examine the validity of FINDR, we generated simulated population spike trains from a known low-dimensional dynamical system and checked whether FINDR can discover latent dynamics that are similar to the ground truth. The low-dimensional dynamical system we use is inspired by the “*n*-bit flip-flop task” (Sussillo & Barak, 2013). In this task, the system receives transient pulse inputs from *n* different channels and needs to memorize the value of the most recent pulse in each channel. It is known that a dynamical system can use attractors to solve this task and that the attractor structure of the system reflects the statistics of the pulses (Kim et al., 2023). For example, in Figure 2a, we let the system memorize the pulse values from two channels, where the pulse value in channel 1, *c*_1_, and the value in channel 2, *c*_2_, are constrained to satisfy 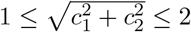. For the system to have robust memory of the pulses in the two channels, it should form a 2-dimensional continuous attractor that has the shape of a disk (Kim et al., 2023).

**Figure 2.**
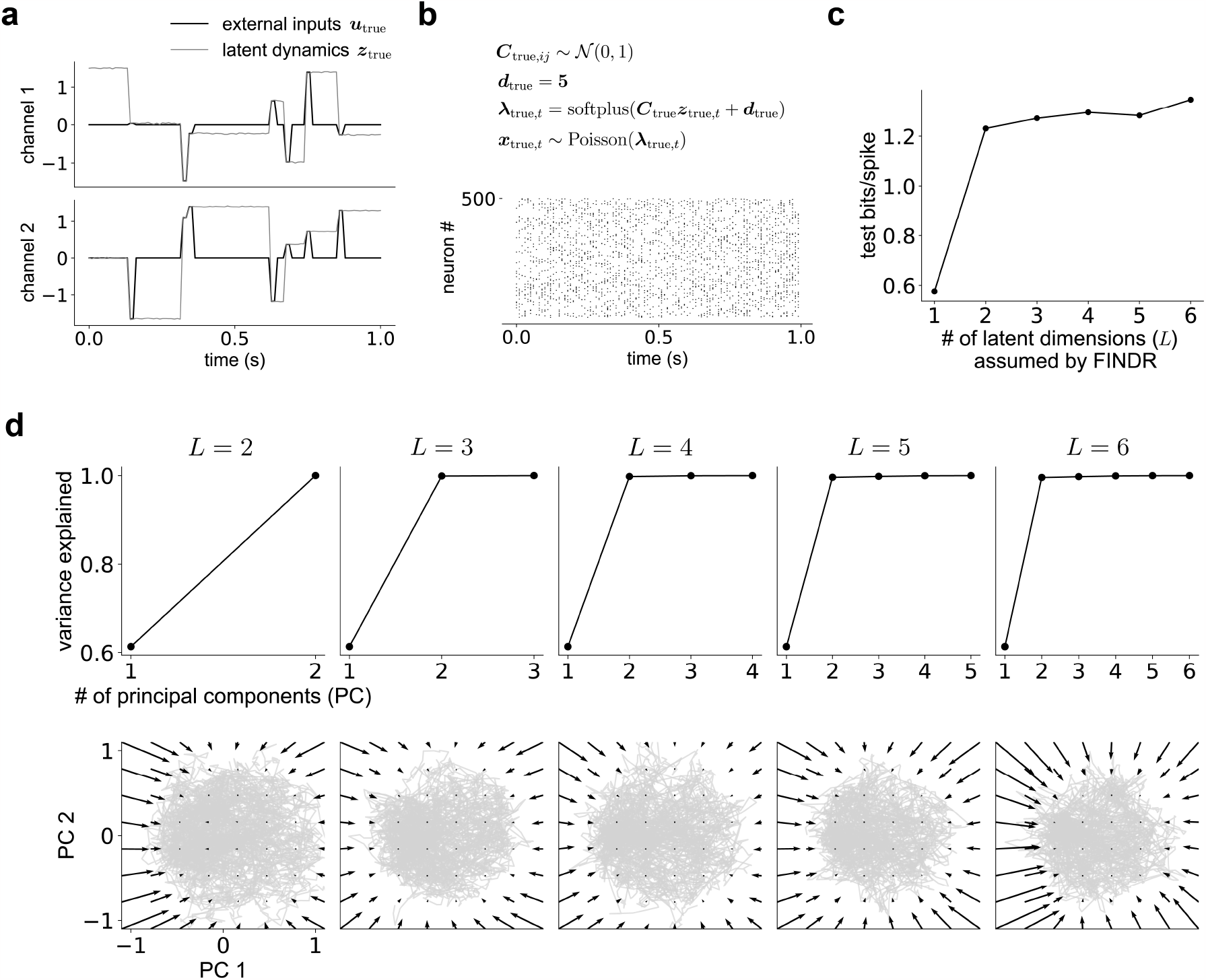
FINDR discovers latent dynamics with the geometry of continuous attractors that reflect how the synthetic neural population stores external inputs. **a**, An example trial with external inputs ***u***_true_ in each channel shown in black, and the state trajectory of the dynamical system that maintains the previous pulse value shown in gray. **b**, An example trial of the simulated population spike trains. **c**, test bits/spike as a function of the latent dimensions (d) assumed by FINDR. **d**, Principal component analysis (PCA) on the inferred latent trajectories across FINDR models assuming *L* = 2, 3, …, 6 show that the first two PCs are sufficient to capture most of the variance in the latent trajectories. The inferred vector fields projected onto the first two PCs are also consistent across FINDR models assuming *L* = 2, 3, …, 6. Gray lines represent latent trajectories.

We simulated our data from 500 different Poisson spiking neurons, with average firing rates around 5 spikes/s (Figure 2b), from the latent trajectories ***z***_true_ that trace, with some small noise, the optimal solution to the task, given external inputs ***u***_true_ (Figure 2a). Then we asked whether FINDR, given the neural population activity and external inputs ***u***_true_, can reconstruct a 2-dimensional disk attractor (Figure 2c-d). To identify whether FINDR correctly captures the true latent dimensionality (*L* = 2) of the population spike trains, we trained multiple FINDR models, each assuming different latent dimensions (*L* = 1, 2, …, 6), on 600 trials of the simulated population spike trains. For each of the FINDR models assuming different latent dimensions, we did a grid search over the hyperparameters (see Section 4.1.6 for details) and found the best-performing model by evaluating bits/spike, which is a normalized log-likelihood score (Pei et al., 2022), on 200 validation trials not used during training. We find that bits/spike, evaluated on 200 test trials (which are separate from the validation trials), saturates around *L* = 2 (Figure 2c). Consistent with this result, we also find that when we do principal component analysis (PCA) on the FINDR-inferred latent trajectories ***z***, we see that two principal components (PCs) are sufficient to explain more than 99% of the variance in each model that assumes a different *L* (= 2, 3, …, 6) (Figure 2d). Furthermore, when we project the vector field inferred from FINDR onto the first two PCs, we find an approximate disk attractor across FINDR models assuming *L* = 2, 3, …, 6 (Figure 2d).

When we do similar analyses with simulated population spike trains generated from a 2-dimensional dynamical system with 4 discrete attractors and a 2-dimensional system with a continuous attractor that has the shape of a rectangle, we find that FINDR discovers these structures (Extended Data Figure 1). Furthermore, for the dynamical system with a rectangular attractor, the width and length of the rectangle were roughly preserved in the inferred latent representation (Extended Data Figure 1c). We find similar results for a 3-dimensional dynamical system with a continuous attractor of the rectangular prism shape (Extended Data Figure 1d-g). Therefore, distance in the FINDR latent space is meaningful. These results suggest that FINDR can infer latent dynamics with attractors of various geometries, and can correctly identify the dimensionality of the latent dynamics.

### 2.5. FINDR outperforms existing methods in capturing real neural population responses

To evaluate FINDR’s performance relative to existing methods in reconstructing neural population activity, we applied FINDR, switching linear dynamical systems model (SLDS; Linderman et al. (2017)), recurrent switching linear dynamical systems model (rSLDS; Linderman et al. (2017)), and autoLFADS (Keshtkaran et al., 2022; Sedler & Pandarinath, 2023) to a dataset comprising 67 simultaneously recorded choice-selective neurons from dorsomedial frontal cortex (dmFC) and medial prefrontal cortex (mPFC) of a rat engaged in a decision-making task for a total of 448 trials (Luo et al., 2023). On each trial, the rat listens to two simultaneous, randomly timed auditory click trains played from loudspeakers on its left and right. At the end of the stimulus, it turns to the side that had the greater total number of clicks for water reward. The spike trains and the auditory click times given to the models were aligned to the stimulus onset (Figure 3a; for a more detailed description of the task and the selection criteria for neurons, see Luo et al. (2023)).

**Figure 3.**
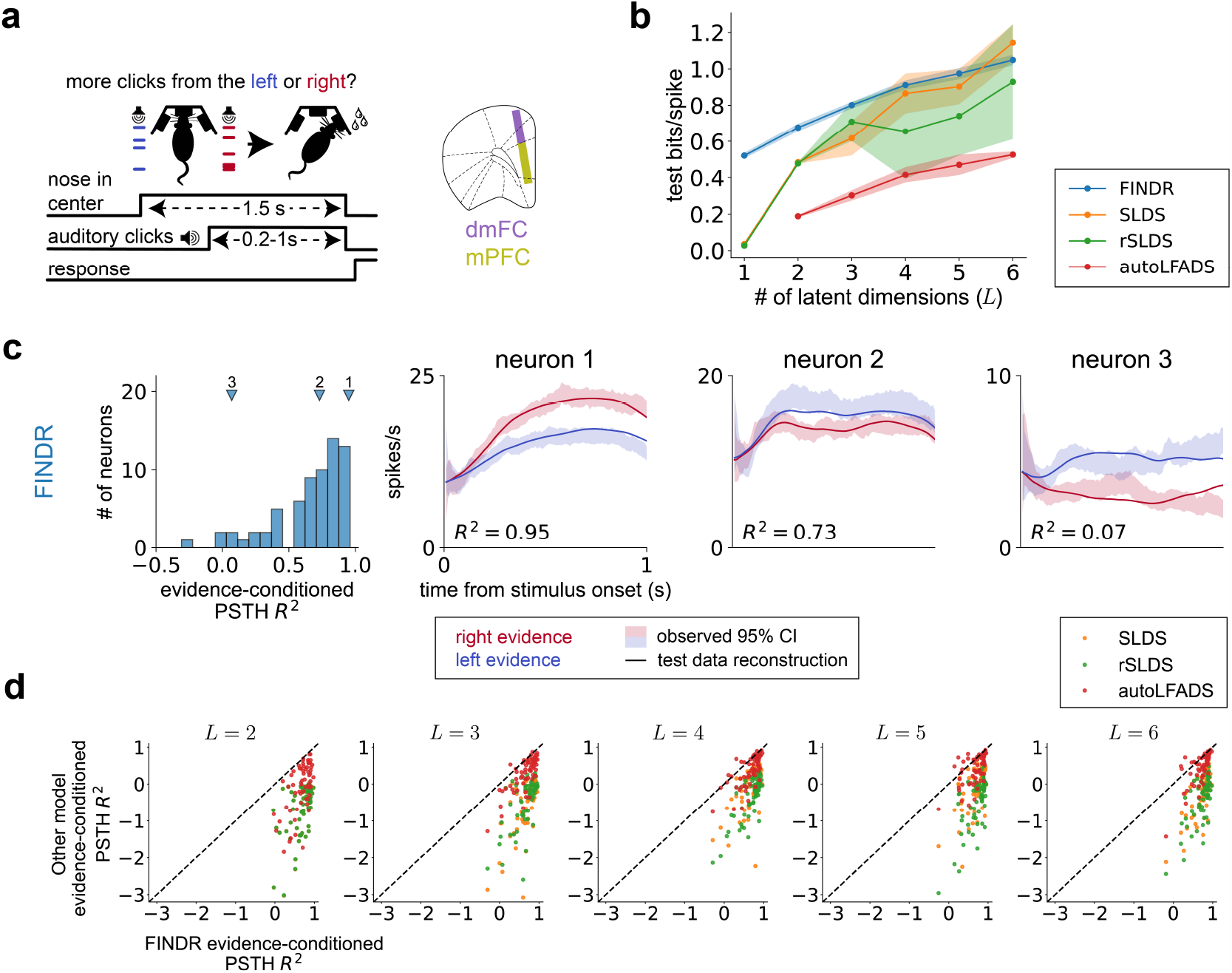
FINDR outperforms SLDS, rSLDS, and autoLFADS in reconstructing the neural population activity of a rat performing perceptual decision-making. **a**, Neurons from dorsomedial frontal cortex (dmFC) and medial prefrontal cortex (mPFC) were recorded while the rat listened to a stream of click trains from its left and right sides of the operant chamber, and oriented to the side that had more clicks to receive water reward. **b**, 5-fold cross-validated bits/spike across different latent dimensions for FINDR, SLDS, rSLDS, and autoLFADS. The shading indicates *±* 1 standard deviation across folds. For SLDS and rSLDS, we consider only the best-performing model among models assuming the number of discrete latent states = {5, 10, 15, 20}. For autoLFADS, *L* corresponds to the factor dimension, not the size of the generator RNN. AutoLFADS with *L* = 1 fails to train. **c**, FINDR with *L* = 3 captures the complex trial-averaged temporal profiles of individual neurons in mPFC and dmFC. The goodness-of-fit is measured using the *R*^2^. **d**, We perform an analysis similar to **c** for other models and compare the *R*^2^ obtained from these models and the *R*^2^ from FINDR.

When we assess the 5-fold cross-validated bits/spike across various models assuming latent dimensions from *L* = 1 to *L* = 6, we find that the bits/spike for FINDR is consistently higher than existing models across dimensions, especially when *L <* 3 (Figure 3b). We also use the match between observed and model-constructed peristimulus time histograms (PSTHs) of individual neurons to measure goodness of fit. Figure 3c shows example neurons’ activity averaged across trials sorted by whether the stimulus favors a leftward or a rightward choice (evidence-conditioned PSTH), and the reconstruction of this conditioned PSTH from FINDR with *L* = 3. The coefficient of determination (*R*^2^) between the observed PSTH and FINDR’s reconstruction is computed for each neuron, and shown as a histogram, with indicators on where the example neurons sit in this distribution (Figure 3c). We repeat this procedure for SLDS, rSLDS, and autoLFADS across different latent dimensions, and find that FINDR consistently outperforms competing models under this metric (Figure 3d, Extended Data Figure 2). These results together suggest that FINDR can reconstruct neural population activity from latent dynamics with fewer dimensions compared to existing methods.

### 2.6. FINDR reveals attractor dynamics in the rat frontal cortical activity during decision-making

That FINDR can capture neural population activity using a latent representation with a lower number of dimensions compared to existing methods allows us to inspect the nature of the learned representation in an intuitive and interpretable manner. In addition to reconstructing latent trajectories from neural population activity (due to the first objective in Figure 1a), FINDR also learns the velocity vector field of the learned latent representation (due to the second objective in Figure 1b-c). In Figure 4a, we explicitly visualize the vector field and the attractor structure discovered by FINDR with *L* = 3 in the first two PC directions for the neural population dataset in Figure 3a. We see that on average, the latent state falls into the attractor in the upper right when the stimulus favors rightward choice and that it falls into the attractor in the bottom left when the stimulus favors leftward choice. Therefore, the inferred latent trajectories and the vector field represent computations that are relevant to the task that the animal performs. We see that the first two PCs explain 95% of the variance in the inferred latent trajectories across FINDR models that assume different latent dimensions (*L* = 2, 3, …, 6). When we project the inferred latent trajectories to the first two PCs, we also see consistency across dimensions (Figure 4b). This suggests that FINDR can discover interpretable task-relevant latent dynamics.

**Figure 4.**
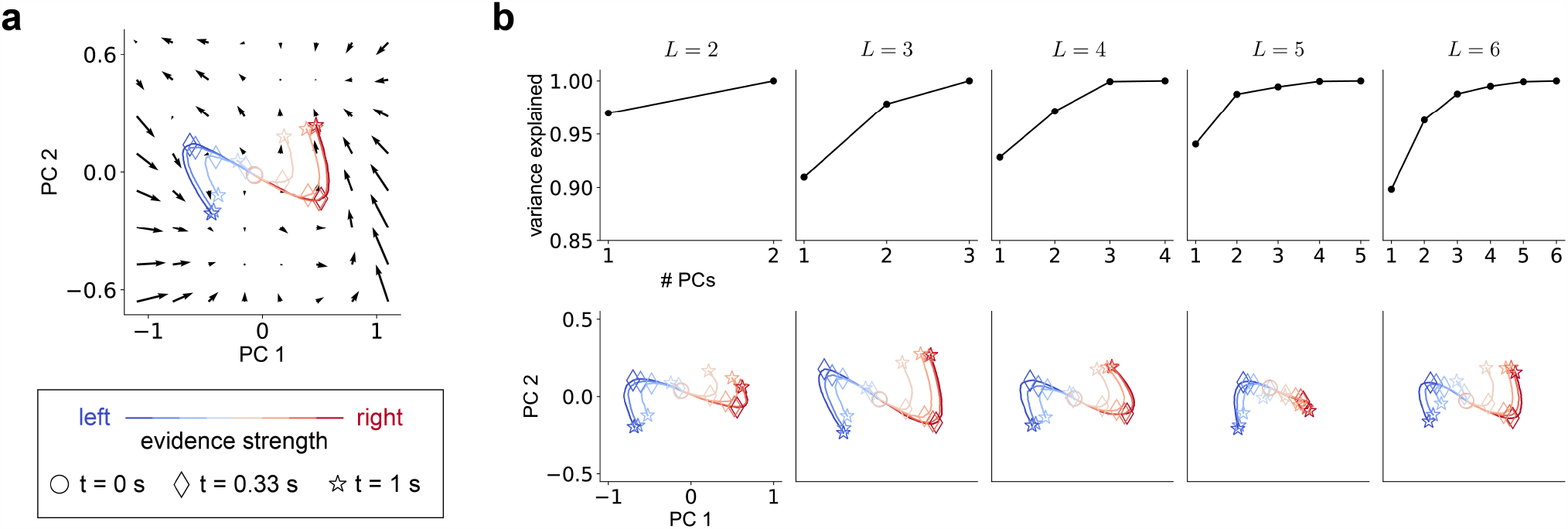
FINDR reveals attractor structure in the latent dynamics of frontal cortex during perceptual decision-making. **a**, FINDR-inferred vector field projected onto the first two principal components (PCs) shows attractor structure. To show how the decision process depends on the input, trials are sorted by their evidence strength, and we show the trial-averaged trajectories as colored lines. **b**, For FINDR models with different latent dimensions, more than 95% of the variance is captured by the first two PCs for *L* = 3, 4, 5, 6. For models with two or more dimensions, the trajectories projected onto the first two dimensions were qualitatively similar.

## 3. Discussion

We introduced an unsupervised deep learning method called FINDR, which infers low-dimensional latent stochastic dynamics underlying neural population activity. When FINDR is trained to spike trains simulated from a system hand-built to exhibit continuous attractors, we demonstrated that FINDR can correctly capture the low-dimensional velocity vector fields and the attractor structure. In a real neurophysiological dataset where the ground truth is not known, we demonstrated how we can increase the interpretability of the latent dynamics discovered by FINDR through separate learning of task-relevant and -irrelevant components in the neural population activity. To validate how well FINDR captures neural activities in this dataset, we demonstrated that FINDR-reconstructed PSTHs of individual neurons match the observed PSTHs across different task conditions. As a comparison, we fit SLDS, rSLDS, and autoLFADS on the same dataset and demonstrated that FINDR outperforms these methods, especially in the regime of low latent dimensions. In addition to strong performance on neural activity reconstruction, FINDR discovered interpretable latent vector fields. As an example, we showed that the rat frontal cortical neurons form attractor dynamics relevant to decision-making (see (Luo et al., 2023) for scientific implications). We plan to show the applicability of FINDR on other neural datasets in the near future.

While we expect FINDR to be generally applicable to a broad range of neural population data, FINDR may be less applicable to certain datasets than others. FINDR is a deep learning-based model that works well with datasets with a high number of simultaneously recorded neurons and trials. While the exact neuron and trial count that give good performance may vary depending on the dynamics in the dataset and the firing rates, generally an increase in the number of neurons should make FINDR’s estimate of each single-trial dynamical trajectory more accurate, while an increase in the number of trials should make FINDR’s estimate of the vector field more accurate, because FINDR has more latent trajectories that traverse the latent space to infer the vector field from. If we ignore the softplus rectification applied to prevent negative firing rates, FINDR uses a linear map to project the dynamics from the latent space to the neural firing rate space (Equation (1)). This means that in datasets where the latent dynamics live in a highly curved manifold instead of a linear subspace of the neural firing rate space, FINDR may have more difficulty fitting the data. If we replace the linear map in FINDR with a nonlinear map, the firing rate predictions could improve. However, it becomes harder to interpret the latent dynamics learned by FINDR. How to learn a latent representation that is interpretable – for example, the one that preserves the geometry of the observables – is an active area of research (e.g., Arvanitidis et al. (2018); Bronstein et al. (2021); Versteeg et al. (2023)), and future investigations are needed to address these challenges.

In conclusion, FINDR extracts low-dimensional dynamical representation from neural population activity. This representation may provide insights into the neural system, spanning multiple levels of descriptions. On one end, FINDR may add a useful constraint when building a biologically plausible network model, as the dynamics of the network should be consistent with the one discovered by FINDR and therefore real neurophysiological data. On the other end, FINDR may facilitate the development of parsimonious, algorithmic-level models of neural computation (see MMDDM in Luo et al. (2023) for an example). These features of FINDR make it a promising approach that can help bridge the gap between the neuronal-level mechanistic description and the algorithmic description of neural function.

## 4. Methods

### 4.1. Model Specification

Dynamics in a neural population may be associated with not only the task that the animal performs but also other factors that are not relevant to the task. Therefore, in FINDR, we distinguish between the task-irrelevant dynamics ***d***(*t*) and task-relevant low-dimensional dynamics ***z***(*t*) and use a separate inference procedure for the two. These latent variables and task stimulus ***u***(*t*) generate spike trains ***y***(*t*) from *N* recorded neurons. We assume that the total length of a trial is *𝒯*, and we bin *𝒯* into *T* bins, with each of the bins having the same width Δ*t*. We assume that there are a total of *M* trials in an experimental session. We denote ***u***_*m,k*_ = ***u***_*m*_(*k*Δ*t*), ***z***_*k,m*_ = ***z***_*m*_(*k*Δ*t*), ***d***_*m,k*_ = ***d***_*m*_(*k*Δ*t*) and ***y***_*m,k*_ = ***y***_*m*_(*k*Δ*t*), with *k* ∈{1, …, *T*} representing the *k*-th time bin within a trial, and *m* ∈ {1, …, M} representing the *m*-th trial. In all tasks that we consider in this work, *𝒯*= 1 second, with Δ*t* = 0.01 second, except for in Section 2.6, where we only considered the epoch of a trial from stimulus onset to right before movement initiation to aid FINDR in learning interpretable representation. Therefore, the length of each trial was variable in Section 2.6. For Section 2.5, for each trial, we looked at the first 1 second from stimulus onset. Thus, if the stimulus duration was less than 1 second on a given trial, we also considered spiking activity during movement. We fixed the lengths of all trials to be the same in this Section to allow comparisons to existing models (datasets with variable trial lengths are not yet supported in some model implementations we consider).

#### 4.1.1. Inference of task-irrelevant dynamics

To model task-irrelevant fluctuations in an individual neuron’s firing rate *across* a total of *M* trials, we compute each neuron *n*’s average firing rate for each trial *m* with

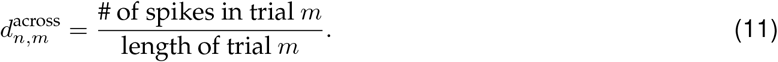

We let the *n*-th element of the *N* -dimensional vector 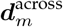 to be 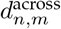. We partition *M* randomized trials into 5 equal subsets, where 3*/*5 of the *M* trials are used for training, 1*/*5 for validation, and 1*/*5 for testing. For each neuron *n*, we fit a linear basis function model (Bishop, 2007) with *p*_across_ radial basis functions to the set of points 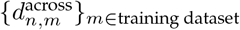. The radial basis functions are evaluated every 1s and for a total of 10, 000s (assuming that no session runs more than 10, 000s). We used the time stamp of the onset of trial *m* (which lies between the beginning of the session, 0s, and 10, 000s, and rounded to the nearest second) as the input to the radial basis functions, which should ideally output 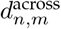 at that time point. We minimize the mean squared error (MSE) to fit. We denote the model’s prediction of 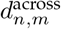 as 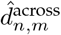. To determine the optimal *p*_across_ (from *p*_across_ ∈ {4, …, 10}), and the optimal *ℓ*_2,across_ regularization coefficient (from *p* ∈ {0.1, 0.1^2^, 0.1^3^, 0.1^4^, 0.1^5^}), we use the validation dataset. We obtain our out-of-sample prediction of 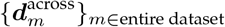using 5-fold cross-validation.

To model task-irrelevant fluctuations in an individual neuron’s firing rate *within* each trial, we again fit, for each neuron *n*, a linear basis function model with *p*_within_ radial basis functions that minimizes the MSE

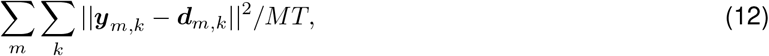

along with an *ℓ*_2,within_ regularization on the weights, where 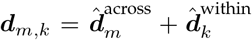 and 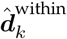 represents the output of the linear basis function model with *p*_within_ radial basis functions. We determine the optimal *p*_within_ and the optimal *ℓ*_2,within_ coefficient using the validation dataset used above, similar to how we determined the optimal *p*_across_ and the optimal *ℓ*_2,across_.

#### 4.1.2. Inference of task-relevant dynamics

For simplicity, going forward we suppress *m* in our notation whenever we can. To maximize the log-likelihood of observing the population spike trains ***y*** given the task-related external inputs ***u***, the task-irrelevant baseline inputs ***d*** and the model parameters *θ*, we need to compute

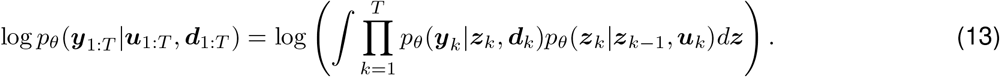

Here, *p*_*θ*_(***y***_*k*_|***z***_*k*_, ***d***_*k*_) is given by a discretized form of Equation (2):

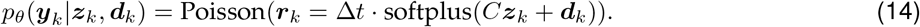

The term *p*_*θ*_(***z***_*k*_|***z***_*k*−1_, ***u***_*k*_) specifies the Euler-Maruyama discretization of the dynamics given by Equation (3):

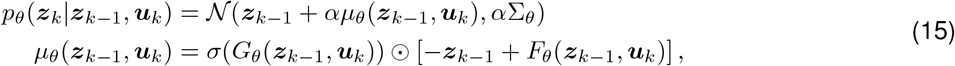

where *α* = Δ*t/τ* . We call Equation (15) the prior process. We assume that the initial condition of the latent ***z***_0_ = **0**. Equation (13) does not have a closed-form solution, and computing this quantity can be computationally expensive. Therefore, we instead compute the evidence lower bound (ELBO) (Kingma & Welling, 2014):

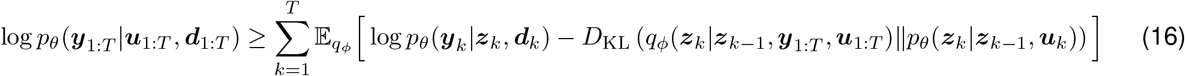

by introducing a variational posterior *q*_*ϕ*_. Here, *D*_KL_ is the Kullback-Leibler divergence between the prior *p*_*θ*_ and the variational posterior *q*_*ϕ*_. We specify *q*_*ϕ*_ as:

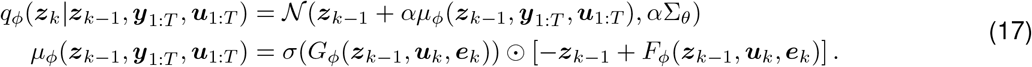

We call Equation (17) the posterior process. The initial condition ***z***_0_ is again set to be **0** here. We also let the diagonal matrix Σ_*θ*_ be the same in both the prior and posterior processes to make the KL divergence finite (Li et al., 2020). 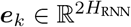 encodes representations of ***y***_1:*T*_ and ***u***_1:*T*_ using a bidirectional GRU (Cho et al., 2014):

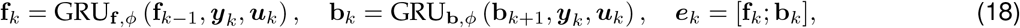

where 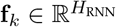 and 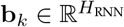. To train FINDR, we compute the gradient of

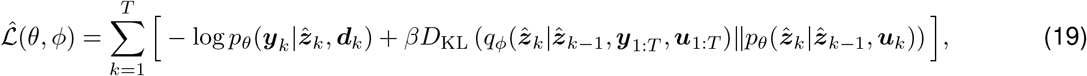

with respect to *θ* and *ϕ* using backpropagation through time (BPTT), where we sampled 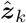 once from *q*_*ϕ*_(***z***_*k*_|***z***_*k*−1_, ***y***_1:*T*_, ***u***_1:*T*_) for *k* ∈ {1, …, *T* }.

With the *m* included back into our notation, Equation (19) can be re-written as

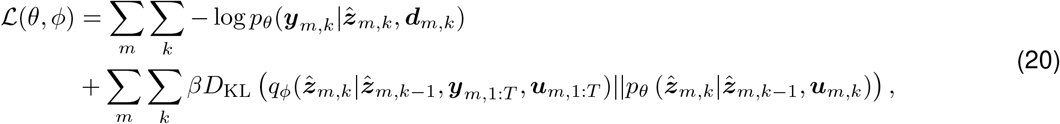

where

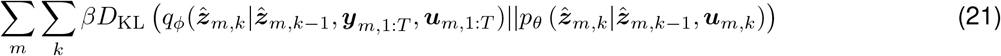

is equal to:

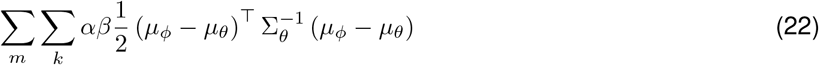

where the arguments of the functions *μ*_*ϕ*_ and *μ*_*θ*_ are 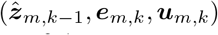 and 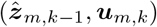, respectively. We set *β* = 2, and thus *c* in Equation (10) is *c* = *αβ/*2 = *α* = 0.1.

#### 4.1.3. Model architecture

For *G*_*θ*_ and *F*_*θ*_ in Equation (8), we use

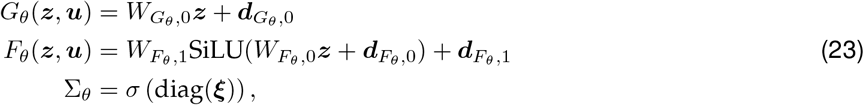

Similarly, for *G*_*ϕ*_ and *F*_*ϕ*_ in Equation (7), we use

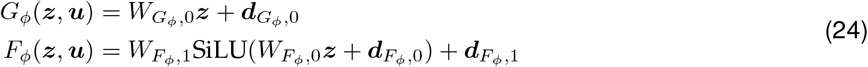

where 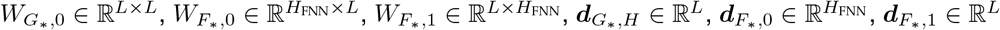and ***ξ*** ∈ ℝ^*L*^ are trainable parameters with *∈ {*θ, ϕ* }. Here, *H*_FNN_ is a hyperparameter that determines the width of the hidden layer. In Equation (18), **f**_0_ and **b**_*T* +1_ are trainable parameters. We let

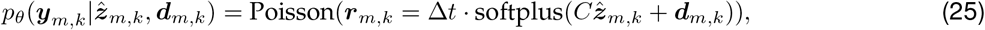

where *C* ∈ ℝ^*N* ×*L*^ is trainable. For all FNNs in this paper, we use the SiLU (a.k.a. swish) activation function (Ramachandran et al., 2017; Elfwing et al., 2017).

#### 4.1.4. Initialization

We use the initialization scheme in Kim et al. (2023) for the kernels that transform the hidden states in our gnSDEs. We use the orthogonal initializer for the kernels that transform the hidden states in the GRUs. We use the Lecun normal initializer (Klambauer et al., 2017) for the kernels that transform the input in both the GRUs and gnSDEs. Biases are i.i.d. normal with variance 10^−6^.

#### 4.1.5. Optimization

To train this model, we use the discrete adjoint sensitivity (i.e., standard backpropagation through time) to compute the gradient of *ℒ* in Equation (20) with respect to {*θ, ϕ*} . A few studies (Gholami et al., 2019; Onken & Ruthotto, 2020) show that the discrete adjoint sensitivity produces more accurate gradients than the continuous adjoint sensitivity used in Chen et al. (2018). We train for a total of 3000 epochs and minimize loss using mini-batch gradient descent with warm restart (Loshchilov & Hutter, 2017). The learning rate increases from 0 to *η* linearly for 10 epochs every 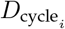 = 2^*i*–1^*D* epochs, where *i* goes from 1 to *i*_end_. After the 10 epochs, the learning rate decays in a cosine manner, where at 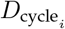, the learning rate becomes 0. *i*_end_ is determined by the minimum 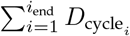 which is greater than or equal to 3000 *D* . is set to be .

#### 4.1.6. Hyperparameter grid-search

For each of the 5-folds in a single experimental session dataset, we do a grid search over *η* = {10^−2.0^, 10^−1.625^, 10^−1.25^, 10^−0.875^, 10^−0.5^ }, *H*_FNN_ = {30, 50, 100 }, and *H*_RNN_ = {50, 100, 200} to identify the model that performs best when the objective is evaluated using the validation dataset. Here, *H*_FNN_ is the number of hidden units in FNNs *F*_*θ*_ and *F*_*ϕ*_, where both networks had a single hidden layer. *H*_RNN_ represents the number of units for both GRU_**f**,*ϕ*_ and GRU_**b**,*ϕ*_. Thus the total number of units for the bidirectional GRU is 2*H*_RNN_.

#### 4.1.7. Fixed hyperparameters

We train FINDR for a total of 3000 epochs. For the first 300 epochs, we train only the first 30 time bins of the trials, and after 300 epochs, we fit all time bins in the trials. We set the coefficient of the *ℓ*_2_ regularization on the weights of all model parameters to be 10^−7^. We let *F*_*θ*_ and *F*_*ϕ*_ be an FNN with a single hidden layer. We set the time constant *τ* = 0.1s. We set *β* = 2. We set the number of trials in a mini-batch to be 25. We set the momentum in mini-batch gradient descent to be 0.9. We perform annealing to the KL term in Equation (21). Specifically, the KL term is multiplied by 1 − 0.99^iteration #^.

### 4.2. Post-Modeling Analysis

While we do not constrain *C* in Equation (25) to be semi-orthogonal (i.e., *C*^⊺^*C* = *I*) during the training procedure, a semi-orthogonal *C* may be desired because distance and angle in the latent space ℝ^*L*^ are distance and angle in the inverse-softplus rate space ℝ^*N*^ . More precisely, ||*C****z*** ^2^|| = ***z***^⊺^*C*^⊺^*C****z*** = ***z***^⊺^***z*** = ||***z***|| ^2^ for all ***z***. Having a semi-orthogonal, and therefore a distance-preserving, map *C* would make the latent trajectories inferred by FINDR more interpretable.

Before we interpret the latent trajectories ***z*** and the inferred velocity vector field *μ*_*θ*_(***z, u***), we perform singular value decomposition (SVD) on *C* = *USV* ^⊺^, where *U* ∈ ℝ^*N* ×*L*^ is a semi-orthogonal matrix, *S* ∈ ℝ^*L*×*L*^ is a diagonal matrix with its entries populated by the singular values and *V*∈ ℝ^*L*×*L*^ is an orthogonal matrix. Then, we apply a transformation 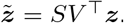. We next perform principal component analysis (PCA) on 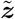 so that the first component of the transformed 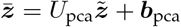corresponds to the first PC, and the *L*-th component of the transformed 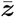corresponds to the *L*-th PC. This transformation by PCA is rigid. Therefore, the distance and angle in the space and axes given by 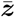are still the distance and angle in the inverse-softplus rate space. The transformed vector field 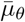 in the space of 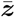is given by:

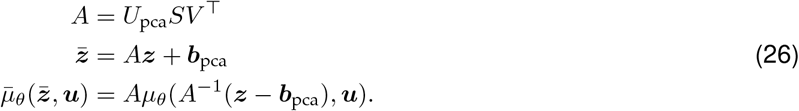

In all of our analyses that show the latent trajectories and vector fields inferred by FINDR, we plot the transformed 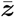 and 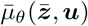. To project the vector field 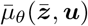 onto the first two PCs, we assume that the third and later components of 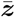 are zero, and consider the first two components of 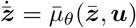.

### 4.3. Fitting Other Models

For SLDS and rSLDS, we use code from https://github.com/lindermanlab/ssm. For autoLFADS, we use code from https://github.com/arsedler9/lfads-torch, with hyperparameter search configurations in configs/pbt.yaml.

## Acknowledgements

We thank Jesse Kaminsky, Chethan Pandarinath, Andrew Sedler, Chris Versteeg, and Iman Wahle for discussions. We thank Jonathan Halverson for help with using the Princeton HPC clusters. This work was supported by grants from NIH R01MH108358, the Simons Collaboration on the Global Brain (SCGB AWD543027), and a U19 NIHNINDS BRAIN Initiative Award (5U19NS104648).

## Author Contributions

T.D.K. conceptualized the method. T.D.K. and T.Z.L. developed the inference method for task-irrelevant dynamics. T.D.K. developed the inference method for task-relevant dynamics. T.Z.L. collected data. T.D.K., T.C., and K.K. developed the gated feedforward neural network (FNN) used in this method. T.D.K. implemented the method as a software package. T.D.K. wrote the manuscript after discussions among all authors. J.W.P. and C.D.B. supervised the project.

---

**Extended Data Figure 1.**
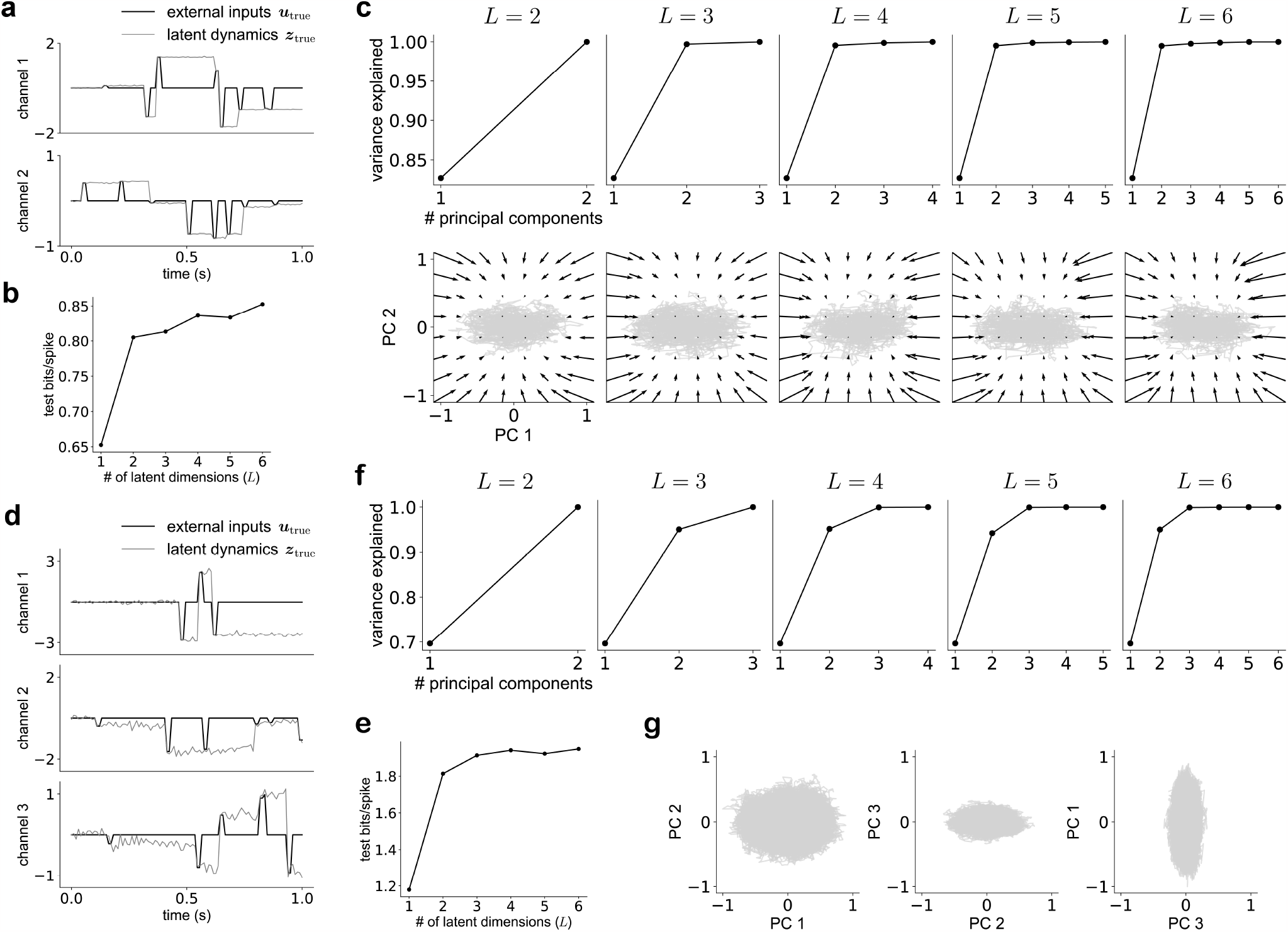
Extended analysis related to Figure 2. **a**, We generate transient pulse inputs from 2 independent channels and let a 2-dimensional system maintain the value of the most recent pulse in each channel. The pulse value in channel 1 (*c*_1_) satisfies − 2≤ *c*_1_≤ 2 and the pulse value in channel 2 (*c*_2_) satisfies − 1≤ *c*_2_ ≤ 1. **b**–**c**, Analysis similar to Figure 2c-d. **d**, We generate transient pulse inputs from 3 independent channels and let a 3-dimensional system maintain (with some noise) the value of the most recent pulse in each channel. The pulse value in channel 1 (*c*_1_) satisfies − 3≤ *c*_1_≤ 3, the pulse value in channel 2 (*c*_2_) satisfies − 2≤ *c*_2_ ≤2, and the pulse value in channel 3 (*c*_3_) satisfies 1 *c*_2_ 1. **e**–**f**, Analysis similar to Figure 2c-d. **g**, The range of values that the latent trajectory ***z*** takes along PC 1 is larger than the range of values that ***z*** takes along PC 2 and PC 3. The range of values that ***z*** takes along PC 2 is larger than the range of values that ***z*** takes along PC 3. This suggests that distance in this latent space is meaningful and reflects the statistics of the pulses.

**Extended Data Figure 2.**
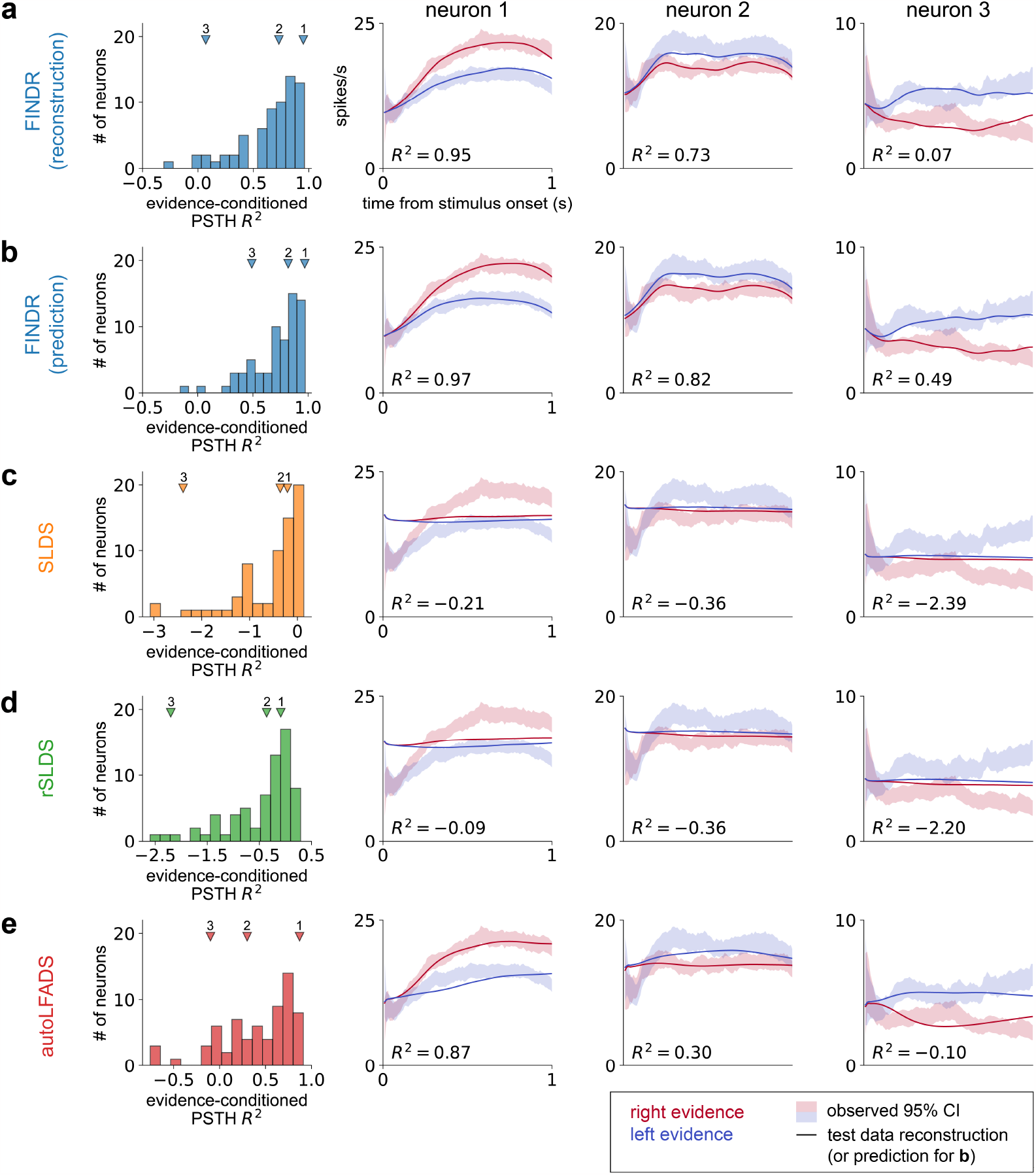
Extended analysis related to Figure 3c. **a**, Same as Figure 3c. **b**, Instead of reconstructing the population spike train ***ŷ***_1:*T*_ from ***y***_1:*T*_ and ***u***_1:*T*_, FINDR can also give a prediction of the population spike train ***ŷ***_1:*T*_ based only on ***u***_1:*T*_ by running FINDR in generative mode. We compute the evidence-conditioned PSTH for each neuron using this prediction. **c**, Analysis similar to Figure 3c, but for SLDS (*L* = 3). **d**, Analysis similar to Figure 3c, but for rSLDS (*L* = 3). **e**, Analysis similar to Figure 3c, but for autoLFADS (*L* = 3).

## References

Arvanitidis, G., Hansen, L. K., and Hauberg, S. Latent space oddity: on the curvature of deep generative models. In International Conference on Learning Representations, 2018.

Bishop, C. M. Pattern Recognition and Machine Learning.Springer, 1 edition, 2007. ISBN 0387310738.

Bronstein, M. M., Bruna, J., Cohen, T., and Veličković, P. Geometric deep learning: Grids, groups, graphs, geodesics, and gauges, 2021.

Burgess, C. P., Higgins, I., Pal, A., Matthey, L., Watters, N., Desjardins, G., and Lerchner, A. Understanding disentangling in β-vae, 2018.

Chen, R. T. Q., Rubanova, Y., Bettencourt, J., and Duvenaud, D. K. Neural ordinary differential equations. In Bengio, S., Wallach, H., Larochelle, H., Grauman, K., Cesa-Bianchi, N., and Garnett, R. (eds.), Advances in Neural Information Processing Systems, volume 31, p .6571–6583. Curran Associates, Inc., 2018.

Cho, K., van Merrienboer, B., Gulcehre, C., Bahdanau, D., Bougares, F., Schwenk, H., and Bengio, Y. Learning phrase representations using rnn encoder-decoder for statistical machine translation, 2014.

Chung, J., Kastner, K., Dinh, L., Goel, K., Courville, A., and Bengio, Y. A recurrent latent variable model for sequential data, 2016.

Cunningham, J. P. and Yu, B. M. Dimensionality reduction for large-scale neural recordings. Nature Neuroscience, 17:1500–1509, 2014.

Driscoll, L., Shenoy, K., and Sussillo, D. Flexible multitask computation in recurrent networks utilizes shared dynamical motifs. bioRxiv, 2022.

Dubreuil, A., Valente, A., Beiran, M., Mastrogiuseppe, F., and Ostojic, S. The role of population structure in computations through neural dynamics. Nature Neuroscience, 25:783–794, 2022.

Duncker, L. and Sahani, M. Dynamics on the manifold: Identifying computational dynamical activity from neural population recordings. Current Opinion in Neurobiology, 70:163–170, 2021.

Duncker, L., Bohner, G., Boussard, J., and Sahani, M. Learning interpretable continuous-time models of latent stochastic dynamical systems. In Chaudhuri, K. and Salakhutdinov, R. (eds.), Proceedings of the 36th International Conference on Machine Learning, volume 97 of Proceedings of Machine Learning Research, pp. 1726–1734. PMLR, 09–15 Jun 2019.

Elfwing, S., Uchibe, E., and Doya, K. Sigmoid-weighted linear units for neural network function approximation in reinforcement learning, 2017.

Gao, Y., Archer, E. W., Paninski, L., and Cunningham, J. P. Linear dynamical neural population models through nonlinear embeddings. In Lee, D., Sugiyama, M., Luxburg, U., Guyon, I., and Garnett, R. (eds.), Advances in Neural Information Processing Systems, volume 29. Curran Associates, Inc., 2016.

Gerstner, W. and van Hemmen, J. L. Associative memory in a network of ‘spiking’ neurons. Network: Computation in Neural Systems, 3(2):139–164, 1992.

Gholami, A., Keutzer, K., and Biros, G. ANODE: Unconditionally Accurate Memory-Efficient Gradients for Neural ODEs. arXiv, 2019.

Higgins, I., Matthey, L., Pal, A., Burgess, C., Glorot, X., Botvinick, M., Mohamed, S., and Lerchner, A. beta-VAE: Learning basic visual concepts with a constrained variational framework. In International Conference on Learning Representations, 2017.

Hopfield, J. J. Neural networks and physical systems with emergent collective computational abilities. Proceedings of the National Academy of Sciences, 79(8):2554–2558, 1982. ISSN 0027-8424. doi: 10.1073/pnas.79.8.2554.

Keshtkaran, M. R., Sedler, A. R., Chowdhury, R. H., Tandon, R., Basrai, D., Nguyen, S. L., Sohn, H., Jazayeri, M., Miller, L. E., and Pandarinath, C. A large-scale neural network training framework for generalized estimation of single-trial population dynamics. Nature Methods, 19:1572–1577, 2022.

Kidger, P., Foster, J., Li, X., and Lyons, T. J. Neural sdes as infinite-dimensional gans. In Proceedings of the 38th International Conference on Machine Learning, volume 139 of Proceedings of Machine Learning Research, pp. 5453–5463.PMLR, 2021a.

Kidger, P., Foster, J., Li, X. C., and Lyons, T. Efficient and accurate gradients for neural sdes. In Advances in Neural Information Processing Systems, volume 34, p .18747–18761. Curran Associates, Inc., 2021b.

Kim, T. D., Luo, T. Z., Pillow, J. W., and Brody, C. D. Inferring latent dynamics underlying neural population activity via neural differential equations. Proceedings of the 38th International Conference on Machine Learning, 2021.

Kim, T. D., Can, T., and Krishnamurthy, K. Trainability, Expressivity and Interpretability in Gated Neural ODEs. Proceedings of the 40th International Conference on Machine Learning, 2023.

Kingma, D. P. and Welling, M. Auto-encoding variational bayes, 2014.

Klambauer, G., Unterthiner, T., Mayr, A., and Hochreiter, S. Self-normalizing neural networks, 2017.

Krishnan, R. G., Shalit, U., and Sontag, D. Structured inference networks for nonlinear state space models, 2016.

Li, X., Wong, T.-K. L., Chen, R. T. Q., and Duvenaud, D. Scalable gradients for stochastic differential equations, 2020.

Linderman, S., Johnson, M., Miller, A., Adams, R., Blei, D., and Paninski, L. Bayesian Learning and Inference in Recurrent Switching Linear Dynamical Systems. In Proceedings of the 20th International Conference on Artificial Intelligence and Statistics, volume 54, p .914–922, 2017.

Loshchilov, I. and Hutter, F. Sgdr: Stochastic gradient descent with warm restarts, 2017.

Luo, T. Z., Kim, T. D., Gupta, D., Bondy, A. G., Kopec, C. D., Elliot, V. A., DePasquale, B., and Brody, C. D. Non-canonical attractor dynamics underlie perceptual decision-making. bioRxiv, 2023.

Macke, J. H., Buesing, L., Cunningham, J. P., Yu, B. M., Shenoy, K. V., and Sahani, M. Empirical models of spiking in neural populations. In Shawe-Taylor, J., Zemel, R., Bartlett, P., Pereira, F., and Weinberger, K. Q. (eds.), Advances in Neural Information Processing Systems, volume 24, p .1350–1358. Curran Associates, Inc., 2011.

Nassar, J., Linderman, S. W., Bugallo, M., and Park, I. M. Tree-structured recurrent switching linear dynamical systems for multi-scale modeling, 2019.

Onken, D. and Ruthotto, L. Discretize-optimize vs. optimize-discretize for time-series regression and continuous normalizing flows. arXiv, 2020.

Pandarinath, C., O’Shea, D. J., Collins, J., et al. Inferring single-trial neural population dynamics using sequential auto-encoders. Nature Methods, 15:805–815, 2018.

Pei, F., Ye, J., Zoltowski, D., Wu, A., Chowdhury, R. H., Sohn, H., O’Doherty, J. E., Shenoy, K. V., Kaufman, M. T., Churchland, M., Jazayeri, M., Miller, L. E., Pillow, J., Park, I. M., Dyer, E. L., and Pandarinath, C. Neural latents benchmark ‘21: Evaluating latent variable models of neural population activity. Advances in Neural Information Processing Systems, 2022.

Ramachandran, P., Zoph, B., and Le, Q. V. Searching for activation functions, 2017.

Rigotti, M., Barak, O., Warden, M. R., Wang, X.-J., Daw, N. D. K. M. E., and Fusi, S. The importance of mixed selectivity in complex cognitive tasks. Nature, 497:585–590, 2013.

Rusch, T. K., Mishra, S., Erichson, N. B., and Mahoney, M. W. Long expressive memory for sequence modeling. arXiv preprint arXiv:2110.04744, 2021.

Sedler, A. R. and Pandarinath, C. lfads-torch: A modular and extensible implementation of latent factor analysis via dynamical systems, 2023.

Sussillo, D. and Barak, O. Opening the black box: low-dimensional dynamics in high-dimensional recurrent neural networks. Neural Computation, 25(3):626–649, 2013.

Versteeg, C., Sedler, A. R., McCart, J. D., and Pandarinath, C. Expressive dynamics models with nonlinear injective readouts enable reliable recovery of latent features from neural activity, 2023.

Vyas, S., Golub, M. D., Sussillo, D., and Shenoy, K. V. Computation through neural population dynamics. Annual Review of Neuroscience, 43:249–275, 2020.

Wang, X.-J. Probabilistic decision making by slow reverberation in cortical circuits. Neuron, 36(5):955–968, 2002.

Yang, G., Joglekar, M., Song, H., Newsome, W., and Wang, X.-J. Task representations in neural networks trained to perform many cognitive tasks. Nature Neuroscience, 22, 2019.

Zoltowski, D., Pillow, J., and Linderman, S. A general recurrent state space framework for modeling neural dynamics during decision-making. In Proceedings of the 37th International Conference on Machine Learning, volume 119, p .11680–11691, 2020.

